# Neuronal expression of Retinoid-Related Orphan Receptor Gamma (RORγ) and revisiting its role in the Central Nervous System

**DOI:** 10.64898/2026.03.03.709291

**Authors:** Logan Reid, Srikar Ganapathiraju, Sophia Mancinelli, Daniel Kagan, Gia Sprouse, Alex Li, Marcello Fazio, William German, Chuangxin Fan, Rojana Saengsot, Paul Feustel, Yunfei Huang

## Abstract

Retinoid-related orphan receptor gamma (RORγ) is a lineage-defining transcription factor for T helper 17 (Th17) cells and has long been considered selectively expressed within the immune system. However, accumulating evidence has implicated Th17 cells and other RORγ-expressing immune populations in embryonic brain development and central nervous system (CNS) function. Notably, RORγ-Cre–mediated deletion of the tuberous sclerosis complex (TSC) has been reported to cause spontaneous seizures and premature lethality, leading to the conclusion that RORγ-expressing immune cells directly regulate CNS physiology. Using multiple RORγ-based reporter mouse lines, we identified unexpected and widespread neuronal labeling throughout the forebrain and cerebellum. RORγ-Cre:GFP^f/^ mice exhibited robust GFP expression in neurons despite the absence of detectable T cells in the brain. Neuronal recombination was evident prenatally and independently validated using the mT/mG reporter line, indicating transient RORγ expression in embryonic neurons. In contrast, a RORγ-GFP reporter showed no detectable RORγ expression in the postnatal brain, either at baseline or following pilocarpine-induced status epilepticus. These findings demonstrate that RORγ expression is not restricted to the immune lineage and reveal previously unrecognized developmental expression in the embryonic brain. Consequently, seizures observed following RORγ-Cre–mediated deletion of TSC1 are unlikely to arise from RORγ-expressing immune cell–dependent mechanisms but instead reflect direct neuronal loss of TSC1. Our results call for careful reinterpretation of prior studies attributing CNS phenotypes to RORγ-expressing immune cells based on RORγ-Cre–driven genetic models.

**Significance:** This work identifies transient neuronal RORγ expression as a critical confounder in immune-targeted genetic studies and reveals a previously unrecognized embryonic expression of RORγ in neurons that may contribute to neurodevelopment. Our findings further raise the possibility that embryonic neurons represent unintended off-targets when deploying therapeutic strategies involving RORγ ligands.

## Introduction

Retinoid-related orphan receptor γ (RORγ) is a member of the retinoic acid receptor– related orphan receptor (ROR) family of nuclear transcription factors, which also includes RORα and RORβ[1]. These receptors regulate gene transcription by binding to ROR response elements (ROREs) within target genes. While RORα and RORβ are broadly expressed in the nervous system and peripheral tissues and play established roles in brain development[1–8], RORγ has been considered largely immune-restricted, with prominent expression in the thymus and secondary lymphoid organs. RORγ is a master regulator of Th17 cells, γδ T cells, and group 3 innate lymphoid cells (ILC3), coordinating IL-17 production and downstream inflammatory programs[9–12]. Consequently, RORγ sits at the center of IL-17 signaling, a pathway critical for host defense, immune homeostasis, and neuroinflammation[13, 14].

IL-17A has been implicated in maternal immune activation (MIA)–induced cortical maldevelopment and behavioral abnormalities in offspring [15], as well as in autism-like phenotypes [16]. Additional studies suggest that Th17 cells influence central nervous system (CNS) function, promoting anxiety- and depression-like behaviors [17–19], while IL-17A overexpression in RORγ transgenic mice alters microglial activity in the dentate gyrus [20]. However, CD4-Cre–mediated deletion of RORγ does not affect anxiety- or depression-like behaviors [21], highlighting ongoing uncertainty regarding the role of RORγ-dependent immune pathways in CNS-associated phenotypes.

In contrast to RORγ, RORβ is essential for cortical layer IV specification and barrel formation[2, 5, 8], and RORβ loss-of-function mutations cause neurodevelopmental disorders and epilepsy in humans [22]. Intriguingly, deletion of TSC1 in RORγ-expressing cells has been reported to induce spontaneous seizures and early mortality [23], suggesting that previously unrecognized RORγ-expressing populations may contribute directly to CNS function. In situ hybridization studies detected RORγ transcripts in the embryonic cortex and cerebellum, with expression peaking around postnatal day 14, although the precise cellular identity of these cells remained unresolved by single-cell RNA sequencing [23].

Under steady-state conditions, T cells are largely excluded from the CNS parenchyma, entering primarily during pathological states [24], and have been identified in resected tissue from patients with epilepsy [25]. In the present study, we employed RORγ-Cre:GFP^f/−^ mice in combination with the mT/mG reporter line to trace RORγ-expressing cells in the brains. Unexpectedly, we observed widespread RORγ-Cre–driven reporter expression in neurons of the cortex, hippocampus, and cerebellum. These findings call for a careful re-evaluation of the role of RORγ in the CNS and the interpretation of RORγ-Cre–based genetic models.

## Materials and Methods

### Animals

Mouse lines including B6.129P2(Cg)-Rorctm2Litt/J (RRID:IMSR_JAX:007572), B6.FVB-Tg(Rorc-cre)1Litt/J (RRID:IMSR_JAX:022791), B6.Cg-Gt(ROSA)26Sortm6(CAG-ZsGreen1)Hze/J (RRID:IMSR_JAX:007906), and B6.129(Cg)-Gt(ROSA)26Sortm4(ACTB-tdTomato,-EGFP)Luo/J (RRID:IMSR_JAX:007676) mice were obtained from The Jackson Laboratory. Both male and female mice were used in all experiments. Animals were housed in a specific pathogen-free, temperature- and humidity-controlled facility under a 12 h light/dark cycle (lights on at 7:00 A.M.) with ad libitum access to food and water. All procedures were conducted in accordance with the guidelines of the Institutional Animal Care and Use Committee and the National Institutes of Health Guide for the Care and Use of Laboratory Animals. Every effort was made to minimize animal suffering and reduce the number of animals used. All methods were performed in accordance with relevant institutional guidelines and regulations.

### Immunohistochemistry and Image Acquisition

Brain harvesting, tissue processing, and immunohistochemistry were performed as previously described [26–28] with minor modifications. Mice were anesthetized with pentobarbital (100 mg/kg, i.p.; Sigma-Aldrich) and transcardially perfused with PBS followed by 4% paraformaldehyde (PFA) in PBS (pH 7.4). Brains were postfixed for 24 hours in 4% PFA and cryoprotected in 30% sucrose in PBS for at least 48 h. Brains were embedded in Neg-50 frozen section medium (*Thermo Fisher Scientific*) and sectioned coronally at 35 μm using a cryostat unless otherwise specified.

For immunostaining, free-floating sections were washed in PBS, then blocked and permeabilized in PBS containing 10% bovine serum albumin (BSA; *Sigma-Aldrich*) and 0.3% Triton X-100 for 2 h at room temperature. Sections were incubated overnight at 4°C with primary antibodies diluted in blocking buffer: anti-GFP (Cat# 132002; *Synaptic Systems*) or anti-NeuN (Cat# MAB377; *Millipore*). After three 5-min washes in PBS, sections were incubated with appropriate fluorescently conjugated secondary antibodies (goat anti-rabbit, Cat# A11034; goat anti-mouse, Cat# A11031; *Thermo Fisher Scientific*) for 2 h at room temperature. Sections were washed three times in PBS, counterstained with DAPI (*Sigma-Aldrich*), and mounted with Fluoromount-G (*Southern Biotech*). Coverslips were sealed with nail polish. Images were acquired using a Zeiss LSM 880 confocal microscope equipped with Airyscan (*Carl Zeiss*) and processed using ZEN Black 2.1 or ZEN Blue Lite 2.3 software (*Carl Zeiss*). Image processing was applied uniformly across experimental groups. GFP and NeuN positive neurons were quantified using Image J.

### Pilocarpine-Induced Status Epilepticus

Status epilepticus (SE) was induced using pilocarpine as previously described [28]. Briefly, 8–10-week-old mice of either sex were injected intraperitoneally with methyl scopolamine (1 mg/kg; *Sigma-Aldrich*) in 0.9% saline 10 min prior to pilocarpine administration. Pilocarpine injections were initiated between 9:00 A.M. and 3:00 P.M. To induce SE, a modified ramping protocol was used: mice received an initial intraperitoneal injection of pilocarpine (200 mg/kg), followed by additional 50 mg/kg injections every 15 min until stage 4 or 5 seizures were observed as previously described [28]. Mice were allowed to remain in SE for 4 h. Afterward, animals were placed on a 30°C warming pad for 1 h to recover.

### Fluoro-Jade B Staining

Fluoro-Jade B (FJB) staining was performed to assess acute excitotoxic neuronal injury following SE. Sections were mounted on 2% gelatin-coated slides and air-dried on a slide warmer at 50°C for at least 30 min. Slides were immersed in 1% sodium hydroxide in 80% ethanol (20 ml of 5% NaOH added to 80 ml absolute ethanol) for 5 min, followed by 2 min in 70% ethanol and 2 min in distilled water. Sections were then incubated in 0.06% potassium permanganate for 10 min on a shaker table and rinsed in distilled water for 2 min. Slides were stained for 20 min in 0.0004% Fluoro-Jade B solution (*Histo-Chem Inc*.) prepared in 0.1% acetic acid. After three 1-min washes in distilled water, excess water was removed by vertical draining (~15 s). Slides were dried on a slide warmer (~50°C) for 5–10 min, cleared in xylene for at least 1 min, and coverslipped with DPX mounting medium (Cat# 06522; *Sigma-Aldrich*).

### Data Analysis

Statistical analyses were performed using GraphPad Prism 10 (*GraphPad Software*). Data are presented as mean ± SEM unless otherwise indicated.

## Results

### RORγ-Cre–mediated expression of GFP reporter in neurons of the adult mouse brain

RORγ-Cre:GFP ^f/−^ mice were used to trace RORγ-lineage cells in the adult mouse brain. Adult mice (~postnatal day 60) were perfused with PBS followed by fixation with 4% PFA. Broad GFP reporter expression was observed in cortical neurons across all layers, with the highest density in layers V and VI (**Fig. 1, Supplemental Fig 1**). In the hippocampus, GFP expression was detected in the pyramidal cell layers and the dentate granule cell layer, with sporadic labeling in the dentate hilus. GFP-positive neurons were also observed in the amygdala (**Supplemental Fig. 1d)**. In the cerebellum, GFP expression was evident in the Purkinje cell layer. A small number of GFP-positive neurons were detected in the basal forebrain (**Supplemental Fig. 1E & 1F**).

**Figure 1.**
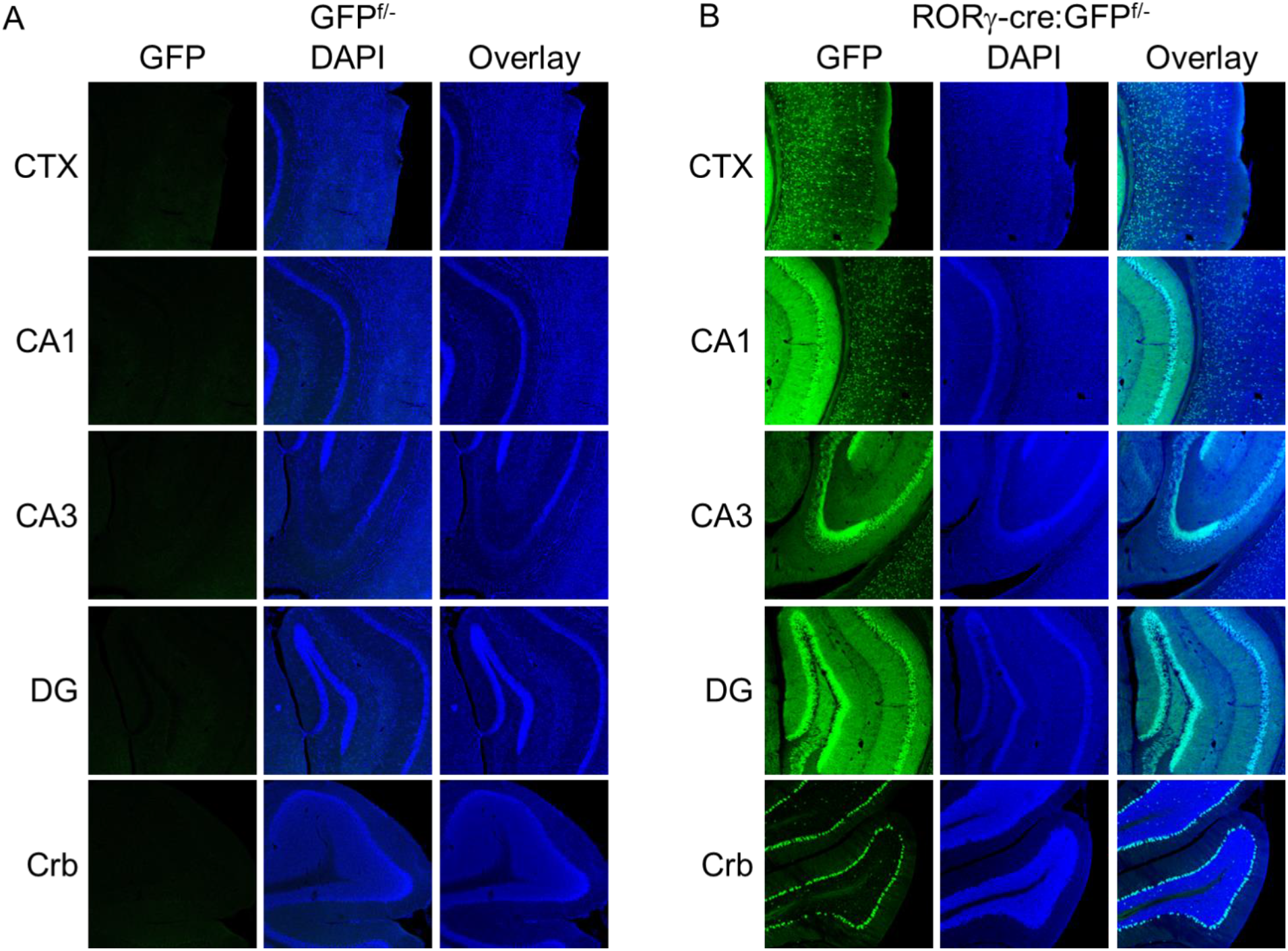
GFP reporter expression in the mouse brain. Representative images from cortex (CTX), hippocampal CA1 and CA3 regions, dentate gyrus (DG), and cerebellum (Crb) of GFP^f/−^ reporter mice (n = 3) (A) and RORγ-Cre:GFP^f/−^ mice (n = 8) (B).

NeuN co-staining revealed that the majority of neurons in the hippocampal pyramidal and dentate granule layers were GFP-positive, as were many neurons in the amygdala (**Fig. 2; Supplemental Fig. 1a**). Approximately 50% of neurons in cortical layers IV and V were GFP-positive, compared with ~15% in layers I–III. In the cerebellum, ~18% of Purkinje cells expressed GFP. GFP^f/−^ mice lacking Cre were used as controls and exhibited only sparse GFP-positive cells, consistent with previously reported low-level leak expression in this reporter line. These findings indicate that the RORγ-Cre line drives reporter expression in a substantial subset of neurons in the adult brain.

**Figure 2.**
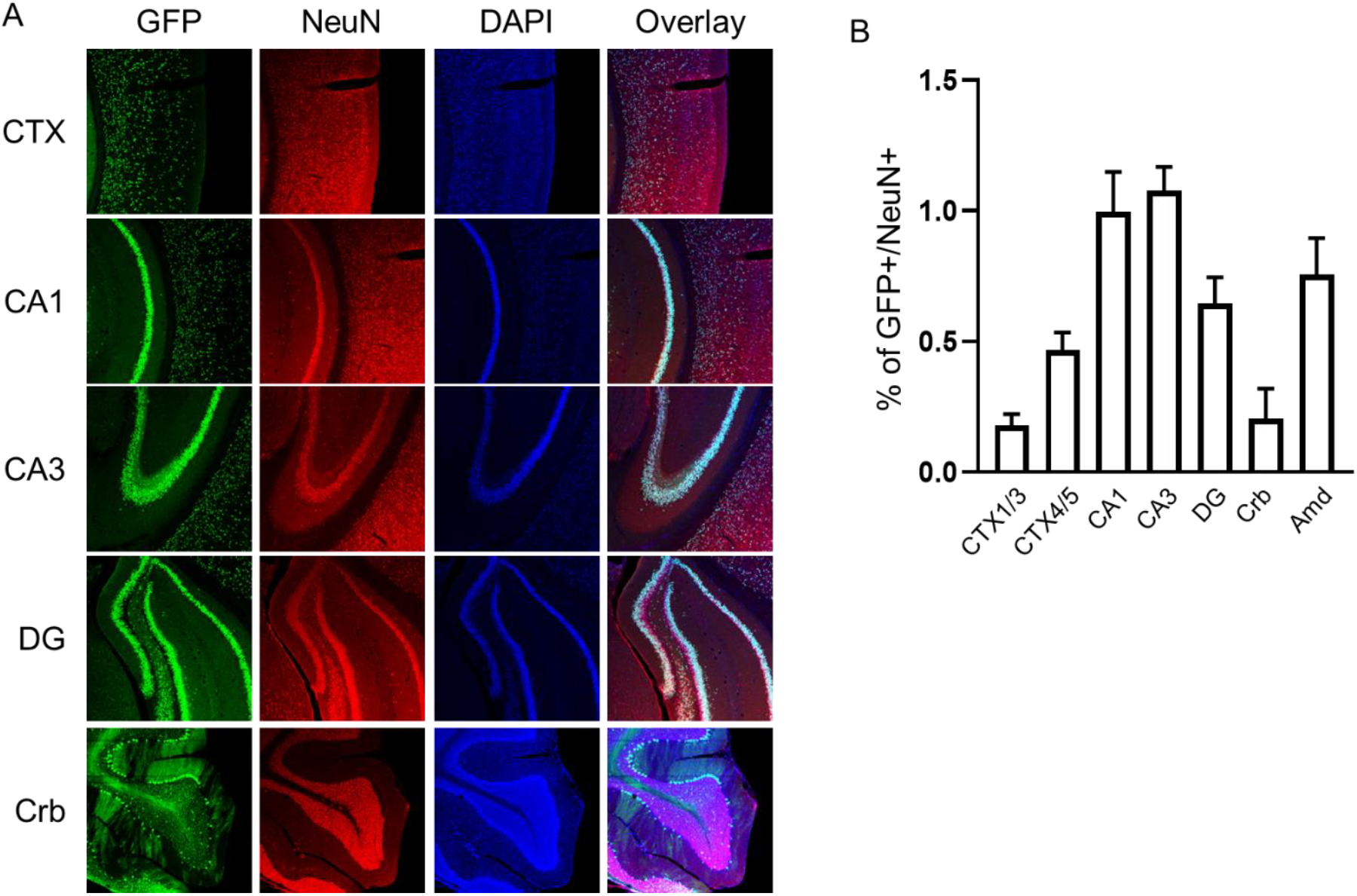
Neuronal localization of GFP reporter expression. (A) Representative images showing colocalization of GFP with NeuN (red) in CTX, CA1, CA3, DG, Crb, and Amd of RORγ-Cre:GFP^f/−^ mice. (B) Quantification of GFP+ cells as a percentage of total NeuN^+ neurons in cortical layers I–III and IV–V, CA1, CA3, DG, Crb, and Amd (n = 4 mice).

### RORγ-Cre–mediated reporter expression during early postnatal development

Given the widespread GFP expression observed in adults, we next examined the onset of reporter expression during postnatal development (**Fig. 3**). GFP expression was detected as early as postnatal day 1 (P1) in cortical and hippocampal neurons. In the cerebellum, sporadic GFP-positive cells were observed at P1 and P3, when cerebellar architecture is still immature. By P7, as cerebellar lamination became more defined, GFP-positive Purkinje cells were clearly evident and aligned along the Purkinje cell layer.

**Figure 3.**
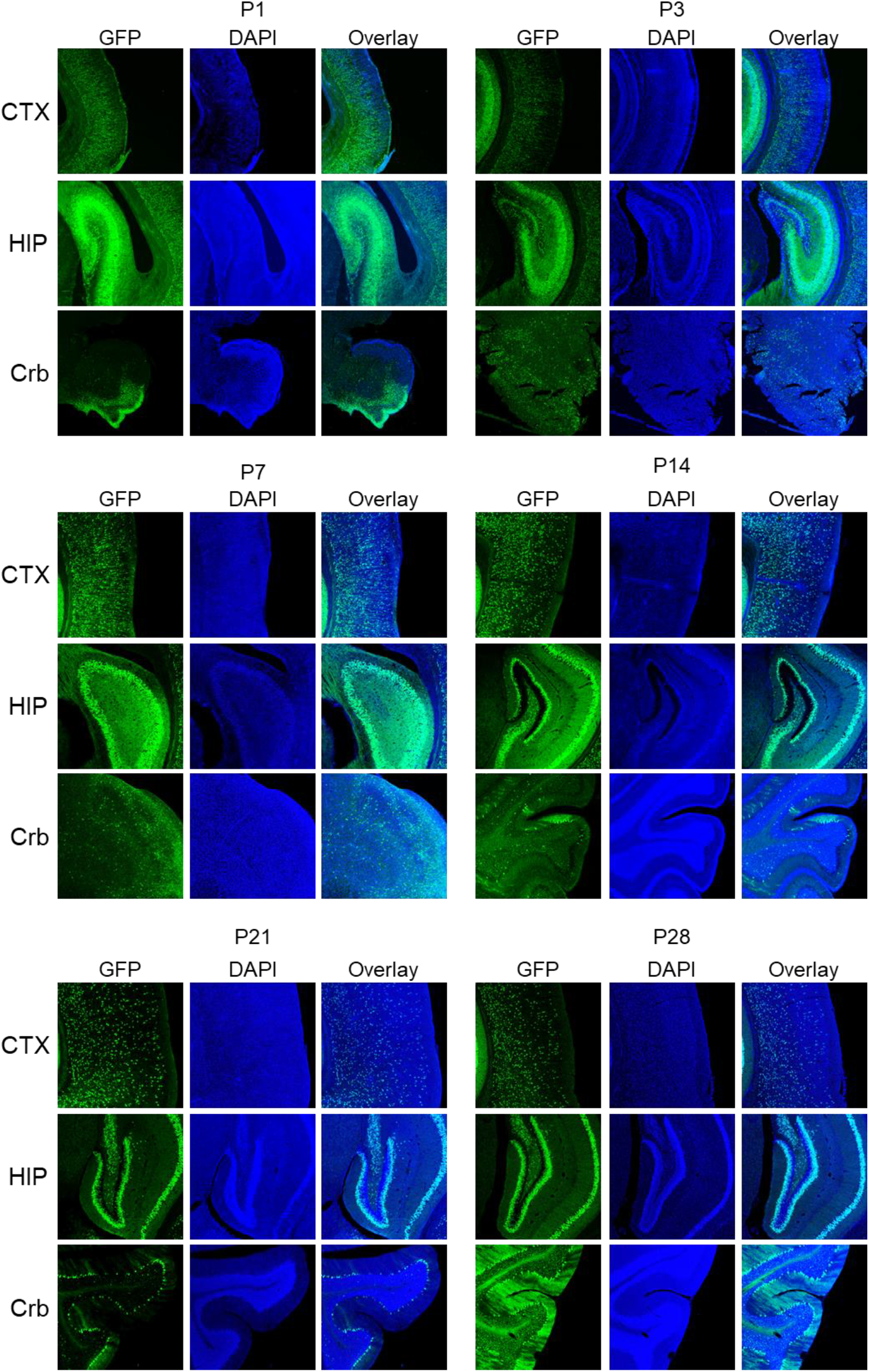
GFP reporter expression during early postnatal development. Representative images of GFP expression in CTX, hippocampus (HIP), and Crb at postnatal day (P)1, P3, P7, P14, P21, and P28 in RORγ-Cre:GFP^f/−^ mice (n = 2–4 mice per time point).

### RORγ-Cre–mediated expression of the mT/mG reporter in postnatal neurons

Because the GFP^f/−^ reporter line exhibits low-level leak expression in cortical and hippocampal neurons [27], we next used the mT/mG reporter line, which is strictly Cre-dependent and does not exhibit detectable leak expression in the CNS [27].

In RORγ-Cre:mT/mG^f/−^ mice, broad membrane GFP expression was observed throughout the cortex, with labeling consistent with membrane-localized GFP. Strong GFP expression was detected in the hippocampal stratum oriens, stratum radiatum, and stratum lacunosum-moleculare near the CA3 region, as well as in the dentate molecular layer and hilus. In the cerebellum, GFP expression was observed in Purkinje cells, with distinct GFP-positive bands across the molecular layer.

Similar to the RORγ-Cre:GFP^f/−^ mice, GFP expression in RORγ-Cre:mT/mG^f/−^ mice was detectable at P3, the earliest examined (**Fig. 4**). In contrast, no GFP expression was observed in mT/mG^f/−^ controls. These findings demonstrate that RORγ-Cre–driven reporter expression is not specific to the GFP^f/−^ line and confirm the presence of RORγ promoter activity during prenatal brain development.

**Figure 4.**
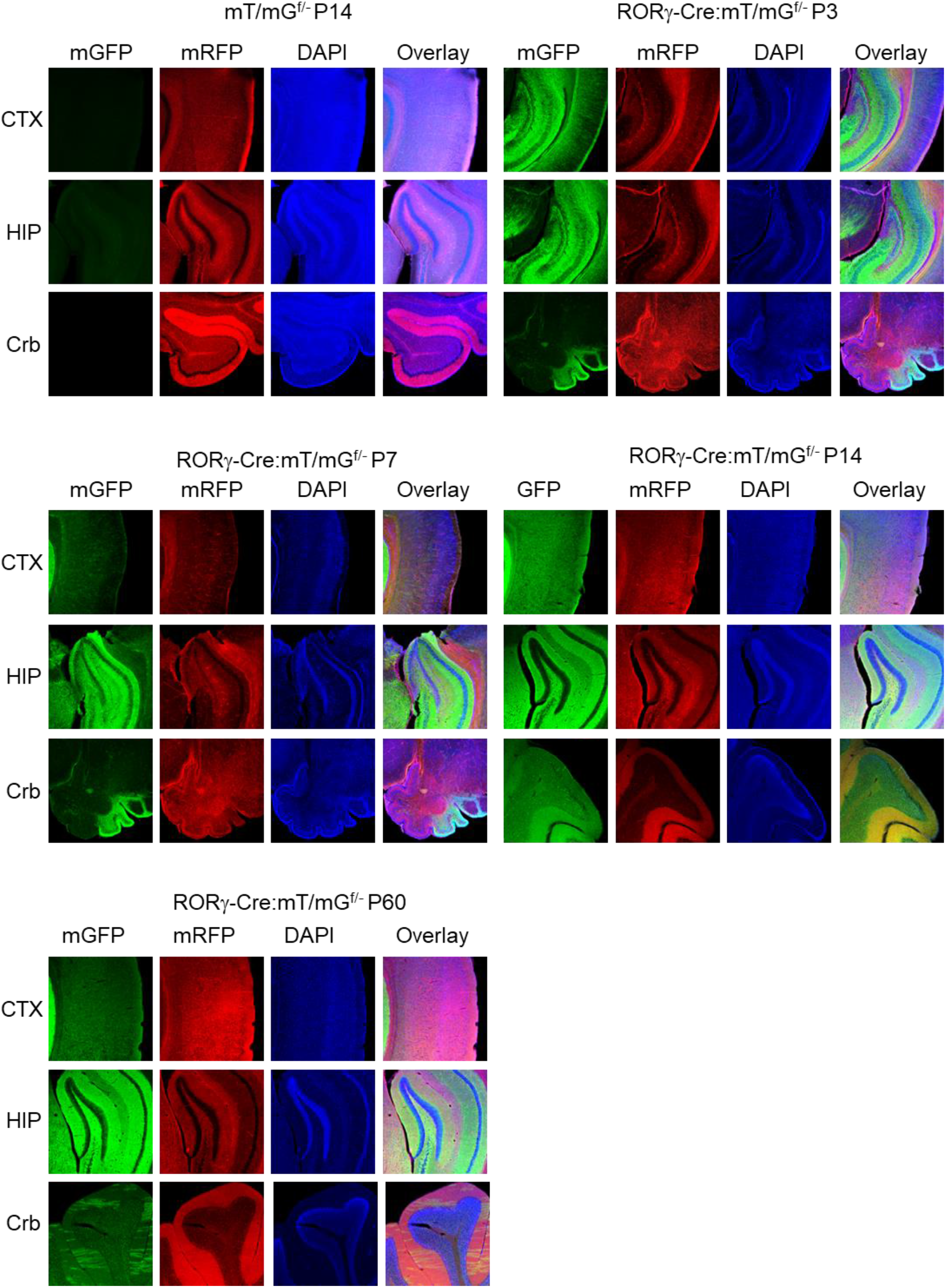
mT/mG reporter expression in RORγ-Cre;mT/mG^f/−^ mice during postnatal development. Representative images of membrane GFP (mG) expression in CTX, HIP, and Crb at P3, P7, P14, and P60 in RORγ-Cre;mT/mG^f/−^ mice (n = 2–4 mice per time point). mT/mG^f/−^ littermates were used as controls.

### Absence of RORγ-GFP expression in postnatal neurons

Previous studies reported peak RORγ mRNA expression at P14 in unidentified CNS cell populations [23]. To directly assess RORγ expression in the postnatal brain, we examined RORγ-GFP reporter mice. No GFP expression was detected at P1, P3, P7, P14, or P21 (**Fig. 5**). To rule out low-level expression, sections were immunostained with anti-GFP antibody; however, no GFP-positive cells were detected. These data indicate that RORγ is not detectably expressed in the postnatal mouse brain.

**Figure 5.**
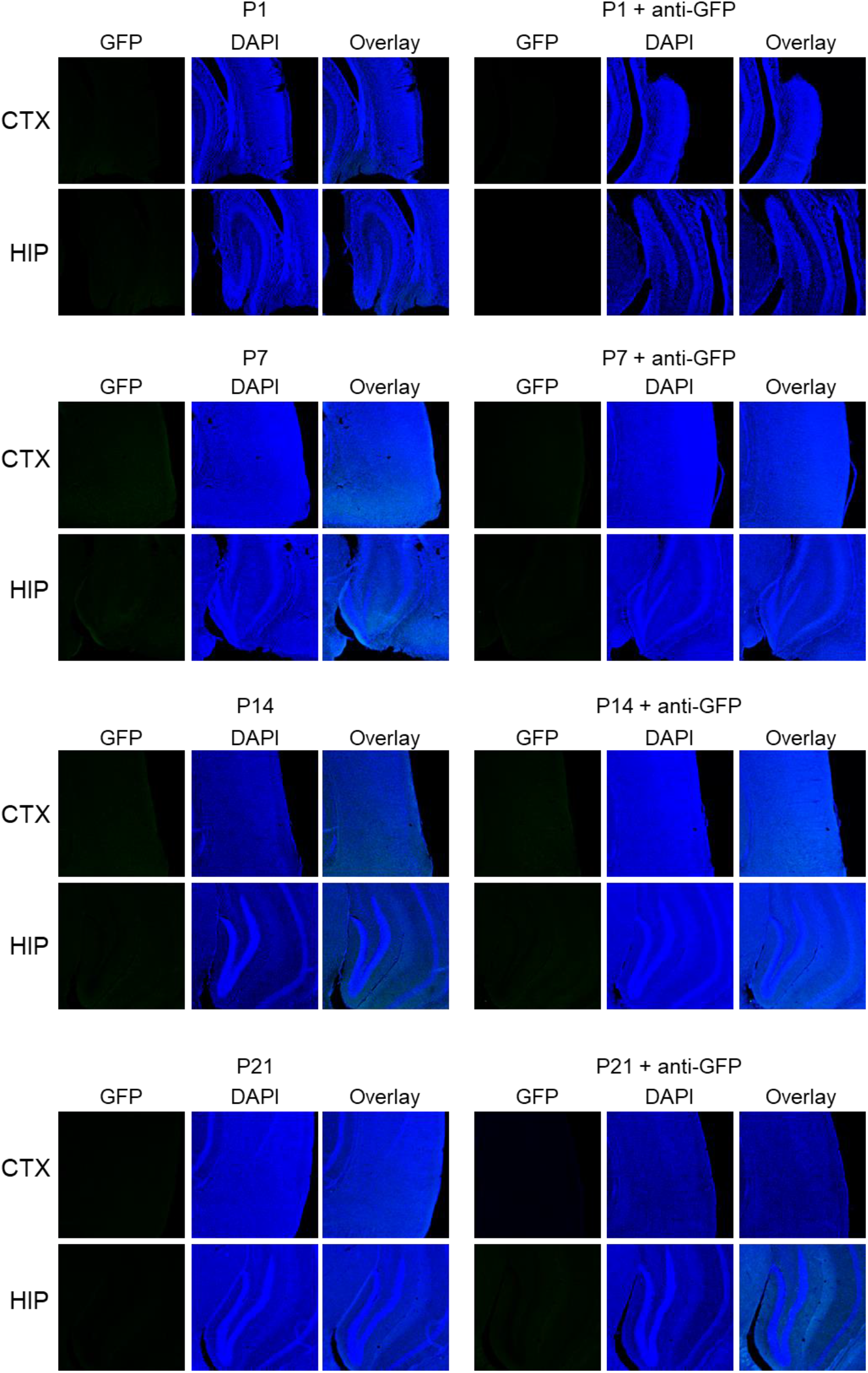
Absence of RORγ-GFP reporter expression during postnatal stages. Representative images showing no detectable GFP expression in CTX and HIP at P1, P7, P14, and P21 in RORγ-GFP mice (n = 2–4 mice per time point). Absence of signal was confirmed by anti-GFP immunostaining.

### Status epilepticus–induced neuronal injury does not induce RORγ-GFP expression

RORγ-Cre–driven reporter activity was prominent in hippocampal neurons, cortical layers IV–V, and the amygdala. We next tested whether neuronal hyperactivity or excitotoxic injury could induce RORγ expression in the CNS. RORγ-GFP mice were subjected to pilocarpine-induced status epilepticus (SE), which causes robust neuronal activation and excitotoxic injury [28]. No GFP expression was detected in the cortex or hippocampus at 3 days post-SE. Fluoro-Jade B staining revealed numerous degenerating neurons in the hippocampal pyramidal layer at post-SE day 3, confirming successful induction of neuronal injury (**Fig. 6**). These results indicate that excitotoxic injury does not induce RORγ expression in postnatal neurons.

**Figure 6.**
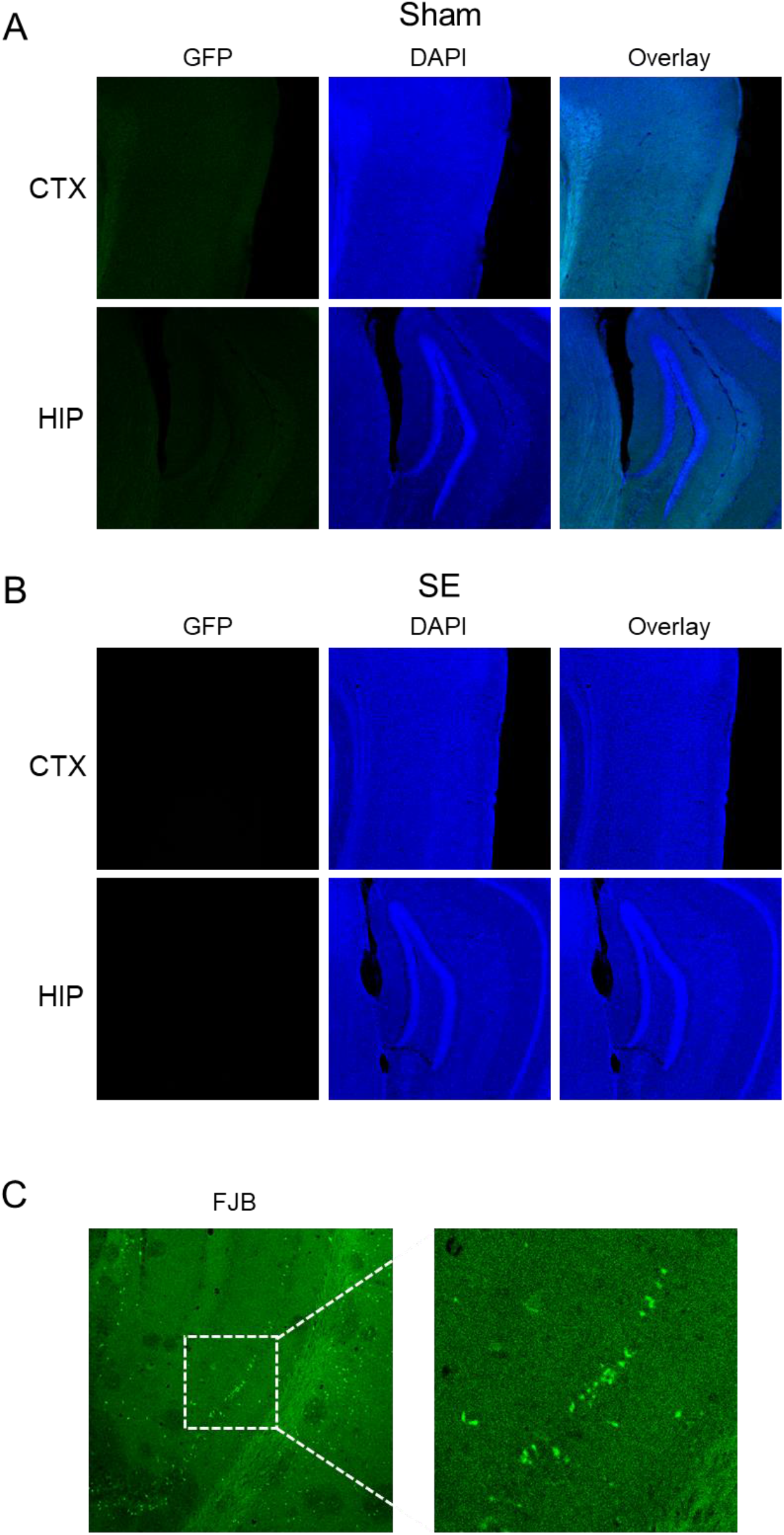
RORγ-GFP reporter expression following status epilepticus. Representative images showing no detectable GFP expression in CTX and HIP of sham-treated (A) and pilocarpine-treated mice (B) following status epilepticus (SE) (n = 4 mice). Fluoro-Jade B staining demonstrates excitotoxic neuronal injury in the hippocampal pyramidal layer after SE.

## Discussion

In the present study, we demonstrate that reporters driven by RORγ-Cre are broadly expressed in neurons of the cortex, hippocampus, and cerebellum. By tracing reporter activity to prenatal stages, we show that RORγ expression occurs during embryonic brain development. Similar to other ROR family members, RORγ is expressed in neurons; however, its expression appears temporally restricted to the embryonic period. A prior report indicated that RORγ mRNA expression peaks at postnatal day 14 [23]. In contrast, we did not detect RORγ-GFP expression at any postnatal stage examined (P3–P21), suggesting that brain expression of RORγ is transient and largely confined to embryogenesis. Although the earlier study did not resolve the cellular identity of RORγ-expressing cells [23], we used two independent Cre-reporter lines to confirm prenatal neuronal expression. While we could not pinpoint the precise embryonic window of RORγ activation, the absence of widespread reporter labeling in the basal forebrain and cortical layers I–III argues against expression in early neural progenitors. Instead, RORγ appears to be expressed in a subset of embryonic neurons, likely including excitatory populations.

The mechanisms governing this transient expression remain unclear. ROR family members are implicated in circadian rhythm regulation [29–32], and both RORβ and RORγ exhibit oscillatory expression patterns [33–35]. Periodic RORγ expression [35] raises the possibility that temporal oscillations contribute to variability in reporter activation among littermates. Although the functional consequences of transient embryonic RORγ expression are unknown, temporally restricted transcriptional activity during critical developmental windows could influence circuit assembly and contribute to inter-individual differences in vulnerability to environmental insults or neurological disease. RORβ regulates transcriptional programs and barrel integrity in cortical layer IV [2, 5, 6, 8], and RORβ loss-of-function mutations cause epilepsy in humans [22]. Whether RORγ similarly shapes cortical development or modulates seizure susceptibility warrants further investigation.

RORγ is a master regulator of Th17 differentiation and IL-17A signaling and has been implicated in neurodevelopmental disorders, including autism spectrum disorders and related psychiatric conditions [15, 16]. In a maternal immune activation (MIA) model, CD4-Cre–mediated deletion of RORγ—presumably in Th17 cells—attenuated MIA-induced cortical abnormalities [15]. However, a subsequent study reported minimal behavioral effects following CD4-Cre–mediated RORγ deletion in adult mice [21], leaving its developmental role unresolved. Our identification of transient embryonic neuronal RORγ expression adds an important dimension to interpreting these findings and suggests that developmental neuronal expression should be considered when evaluating RORγ-dependent phenotypes.

Deletion of TSC1 in neurons, astrocytes, or microglia is well established to cause seizures and premature mortality [26, 36, 37]. Here, we provide strong evidence that the RORγ-positive cells responsible for seizures following TSC1 deletion, as previously reported [23], are neurons rather than immune cells. Accordingly, prior conclusions attributing seizure phenotypes to RORγ-expressing immune populations require reconsideration, and the specific contribution of immune-restricted RORγ deletion to seizure susceptibility remains to be directly tested.

In summary, we identify transient embryonic neuronal expression of RORγ, revealing a previously unrecognized developmental dimension of RORγ biology. These findings open new avenues for investigating the role of RORγ in early brain development and neurodevelopmental disorders. Beyond immunity, RORγ has also been implicated in tumorigenesis [38], and synthetic RORγ agonists and inverse agonists are under active development for autoimmune diseases and cancer [31, 39–42]. Our results raise the possibility that embryonic neurons may represent unintended off-targets of RORγ-targeted therapies, underscoring the importance of accounting for developmental expression patterns in translational strategies.

## Abbreviations

RORα: Retinoid-related orphan receptor alpha
RORβ: Retinoid-related orphan receptor beta
RORγ: Retinoid-related orphan receptor gamma
Th17: T helper 17
CNS: Central nervous system
TSC: The tuberous sclerosis complex
ROREs: ROR response elements
ILC3: Innate lymphoid cells
MIA: Maternal immune activation
SE: Status epilepticus
FJB: Fluoro-Jade B
PBS: Phosphate Buffered Saline
GFP: Green Fluorescent protein
P: postnatal day
CTX: Cortex
HIP: Hippocampus
CA1: Cornu Ammonis area 1
CA3: Cornu Ammonis area 1
DG: Dentate Gyrus
Crb: Cerebellum
Amd: Amygdala
SE: Status epilepticus

## Contributions

L.R., S.G., G.S., C.X.F., and R.S. established mouse colonies and performed animal work. L.R., S.G., S.M. G.S., C.X.F., R.S., D.K., A.L., M.F., and W.G. performed experiments. Y.F.H. conceived and designed the study, prepared the figures, and wrote the manuscript. L.R. and S.M. assisted with manuscript preparation. P.F. contributed to experimental design and provided conceptual guidance. All authors reviewed and approved the final manuscript.

