## Supplemental Figure 1 for "Neuronal expression of Retinoid-Related Orphan Receptor Gamma (RORγ) and revisiting its role in the Central Nervous System"

**Supplementary Figure 1. Forebrain distribution of GFP reporter expression.**
Representative full-montage images of the forebrain from RORγ-Cre:GFP^f/−^ mice (A–C). Higher-magnification images showing GFP expression in the amygdala (Amd) (D) and basal forebrain regions (E, F).

Supplemental Figure 1

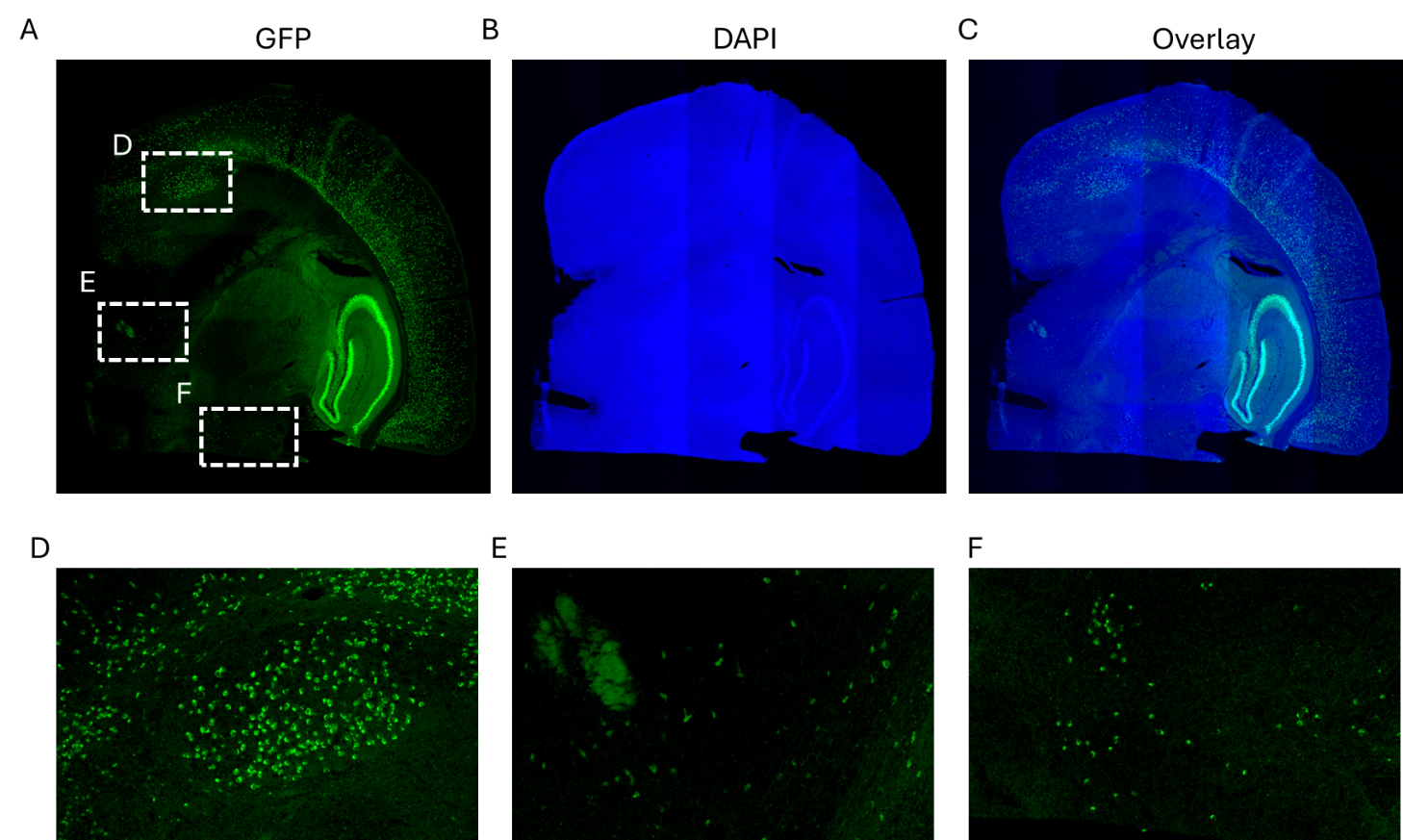
